# Head motion predictability explains activity-dependent suppression of vestibular balance control

**DOI:** 10.1101/560664

**Authors:** H Dietrich, F Heidger, R Schniepp, PR MacNeilage, S Glasauer, M Wuehr

**Affiliations:** German Center for Vertigo and Balance Disorders, University Hospital, LMU Munich, Germany; Department of Neurology, University Hospital, LMU Munich, Germany; Department of Psychology, Cognitive and Brain Sciences, University of Nevada, USA; Institute of Medical Technology, Brandenburg University of Technology Cottbus-Senftenberg, Senftenberg, Germany

**Keywords:** vestibular system, galvanic vestibular stimulation, balance, locomotion, efference copy, feed-forward regulation

## Abstract

Vestibular balance control is dynamically weighted during locomotion. This might result from a selective suppression of vestibular inputs in favor of a feed-forward balance regulation based on locomotor efference copies. The feasibility of such a feed-forward mechanism should however critically depend on the predictability of head movements (PHM) during locomotion. To test this, we studied in healthy subjects the differential impact of a stochastic vestibular stimulation (SVS) on body sway (center-of-pressure, COP) during standing and walking at different speeds using time-frequency analyses and compared it to activity-dependent changes in PHM. SVS-COP coupling decreased from standing to walking and further dropped with faster locomotion. Correspondingly, PHM increased with faster locomotion. Furthermore, SVS-COP coupling depended on the gait-cycle-phase with peaks corresponding to periods of least PHM. These findings support the assumption that during stereotyped human self-motion, locomotor efference copies selectively replace vestibular cues, similar to what was previously observed in animal models.

## 1. Introduction

The vestibular system encodes head orientation and motion to facilitate balance reflexes that ensure postural equilibrium during passive as well as self-initiated movements (Angelaki & Cullen, 2008). During locomotion, i.e., stereotyped self-motion, vestibular influences on balance control appear to be dynamically up- or down-regulated in dependence on the phase and speed of the locomotor pattern. Accordingly, the gain of vestibulospinal reflexes exhibits phasic modulations across the locomotor cycle (Blouin et al., 2011; Dakin et al., 2013) with the result that balance is particularly sensitive to vestibular perturbations at specific phases of the gait cycle (Bent et al., 2004). Furthermore, vestibular influences appear to be down-weighted during faster locomotion. Accordingly, the destabilizing impact of a vestibular loss or perturbation on the gait pattern decreases with increasing locomotion speeds (Brandt et al., 1999; Jahn et al., 2000; Schniepp et al., 2017; Schniepp et al., 2012; Wuehr et al., 2016).

It was previously assumed that activity-dependent modulations of vestibular balance reflexes might reflect an up- or down-regulation of a concurrent intrinsic feed-forward control of posture (Lambert et al., 2012; MacNeilage & Glasauer, 2017; Roy & Cullen, 2004). Accordingly, balance adjustments during self-motion might not solely rely on sensory feedback about how the body has moved, but also on predictions of resultant movements derived from efference copies of the motor command (Straka et al., 2018). Physiological evidence for such a direct feed-forward control mode has recently been shown for animal locomotion. During *Xenopus laevis* tadpole swimming, intrinsic efference copies of the locomotor command deriving from spinal central pattern generators (CPG) were shown to directly trigger ocular adjustments for gaze stabilization and selectively cancel out afferent vestibular inputs (Lambert et al., 2012; von Uckermann et al., 2013). Thus, also during human stereotyped locomotion, efference copies might provide estimates of resultant head motion and assist or even substitute vestibular feedback cues in gaze and balance regulation. The feasibility of such a direct feed-forward mechanism should however critically rely on the predictability or stereotypy of head movements during locomotion (Chagnaud et al., 2012).

Following this intuition, a statistically optimal model was recently proposed, that relates an empirically quantified metric (i.e. the kinematic predictability metric) of head motion predictability to the relative weighting of vestibular vs. motor efference copy cues in gaze and balance regulation during locomotion (MacNeilage & Glasauer, 2017). According to the model, activities linked to less stereotyped head movements should be more dependent on vestibular cues than activities with highly predictable head motion patterns. Likewise, timepoints during the stride cycle when head movement is less stereotyped should exhibit more vestibular dependence. To assess this hypothesis, we examined whether activity-dependent modulations of vestibular balance reflexes can be explained by alterations in the predictability of head movements. Modulations in vestibular balance control during different activities, i.e., standing as well as slow or faster walking, were studied by analyzing the differential impact of a continuous stochastic vestibular stimulation (SVS) on body sway (i.e., center-of-pressure-displacements, COP) in the frequency (coherence) and time (cross-correlation, phase) domain. In parallel, we quantified the predictability of head kinematics associated with these activities and related this metric to an estimate of relative sensory weight.

Using this theoretical and evidence-based approach we aimed to evaluate three hypotheses concerning the role of vestibular cues in balance regulation: (1) Vestibular influence on balance control should decrease from standing to walking due to the presence of a locomotor efference copy; (2) the gain of vestibular balance reflexes should depend on locomotor speed due to increasingly stereotyped head kinematics during faster locomotion; (3) phasic modulations of vestibular balance reflexes across the gait cycle should reflect phase-dependent alterations in head motion predictability.

## 2. Materials and methods

### Subjects

Ten healthy subjects (mean age 29.3 ± 3.7 years, 3 females) participated in the study. None of the participants reported any auditory, vestibular, neurologic, cardio-vascular or orthopedic disorders. All subjects had normal or corrected-to-normal vision. Each participant gave written informed consent prior to the experiments. The local Ethics Committee approved the study protocol, which was conducted in conformity with the Declaration of Helsinki.

### Galvanic vestibular stimulation

A pair of conductive rubber electrodes was attached bilaterally over the left and right mastoid process behind the ears. Stochastic vestibular stimulation (SVS) delivered via this electrode configuration with the head facing forward primarily elicits a postural roll response in the frontal plane (Fitzpatrick & Day, 2004; Schneider et al., 2002). Before electrode placement, the skin surface at the electrode sites was cleaned and dried, and a layer of electrode gel was applied before electrode placing to achieve uniform current density and minimize irritation to the skin during stimulation. The SVS profile consisted of a bandwidth-limited stochastic stimulus (frequency range: 0–25 Hz, peak amplitude ± 4.5 mA, root mean square 1.05 mA) delivered via a constant-current stimulator (Model DS5, Digitimer, Hertfordshire, UK).

### Test procedures

Each participant stood and walked on a pressure-sensitive treadmill (Zebris®, Isny, Germany; h/p/cosmos®, Nussdorf-Traunstein, Germany; 1.6 m long; sampling rate of 100 Hz). Five different conditions were tested in randomized order: three stimulation conditions with continuous SVS and two non-stimulation conditions. SVS was presented during 180 s of quiet standing, as well as during slow walking at 0.4 m/s and medium walking at 0.8 m/s, each for 600 s. Head movements without SVS stimulation were recorded during walking at 0.4 and 0.8 m/s, each for 300 s. Walking was guided by a metronome with a cadence of 52 steps/min for the slow and 78 steps/min for the medium walking speed, respectively. Walking speeds and cadences were chosen in order to allow direct comparison with previous studies (Blouin et al., 2011; Dakin et al., 2013; Iles et al., 2007). During trials, participants were instructed to fixate on a target located 3 m in front of them at eye level. Before each recording, participants were given 30 s to acclimatize to the preset treadmill speed and walking cadence. Between trials, participants were given at least two minutes to recover.

### 2.2 Data analysis

#### Center-of-pressure displacements, head kinematics, and gait parameters

For each stance and walking trial, the continuous trajectory of the center-of-pressure (COP) was computed as the weighted average of the pressure data recorded from the treadmill by using the standard method for determining the barycenter (*sum of mass × position*)/*sum of mass* (Terrier & Deriaz, 2013). COP motion was analyzed in the medio-lateral (ML) dimension, i.e., the primary dimension of postural responses induced by binaural bipolar SVS (Fitzpatrick & Day, 2004). Head kinematics in ML dimension (i.e., linear head acceleration in the ML dimension and angular head velocity in the roll plane) were measured with an inertial measurement unit (IMU) containing a triaxial accelerometer and gyroscope (APDM, Inc., Portland, OR, sampling rate of 128 Hz), strapped to the forehead. Furthermore, for each walking trial, the following spatiotemporal gait parameters were analyzed: base width, stride length, stride time, single support percentage and double support percentage, as well as the coefficient of variation (CV) of each of these parameters.

#### Cross-correlation and coherence analysis

For all stimulation trials, correlation analysis in the frequency (coherence) and time (cross-correlation, phase) domain was used to estimate the average SVS-induced variations in COP-displacements. Coherence estimates with confidence limits were computed based on the auto-spectra of the SVS and COP signals (*P*_*AA*_(*f*) and *P*_*BB*_(*f*) respectively) as well as the cross-spectrum (*P*_*AB*_(*f*)) using a finite fast Fourier transform with a block size of 2 s resulting in a frequency resolution of 0.5 Hz (Rosenberg, Amjad, Breeze, Brillinger, & Halliday, 1989):

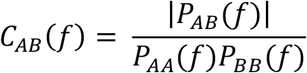

This yielded 95% confidence limits for coherence estimates of 0.033 for stance and 0.010 for walking trials respectively. The resultant coherence estimate is a unitless measure bounded between 1 (indicating a perfect linear relationship) and 0 (indicating independence between the two signals).

Cross-correlations between SVS and COP signals were computed to determine the onset and peak of SVS-induced COP displacements. For this purpose, the inverse Fourier transform of the cross-spectrum *P*_*AB*_(*f*) was computed and normalized by the norm of the input vectors to obtain unitless correlation values bounded between −1 and 1 (Blouin et al., 2011). Resultant 95% confidence limits for cross-correlation estimates were 0.015 for stance and 0.008 for walking trials respectively. Finally, phase estimates between the SVS and COP signals were estimated from the complex valued coherence function. This allows to determine the phase lag corresponding to frequency bandwidths with significant SVS-COP coherence estimates (Dakin et al., 2007). The slope of the phase values over the range of significant coherence estimates was computed using regression analysis and multiplied by 1000/2*π* to yield an estimate of the phase lag in milliseconds.

For the two walking stimulation trials, we further analyzed phasic modulations in the correlation between SVS and COP signals across the average gait cycle, using time-dependent coherence analysis according to a previously described procedure (Blouin et al., 2011; Dakin et al., 2013). First SVS and COP signals were cut into individual strides synchronized to the left heel strike and then time-normalized by resampling each stride to a total of 300 samples. The first 250 strides of each trial were taken for further analysis and padded at the start and end with data from the previous and subsequent strides to avoid distortions in the subsequent correlation analysis. Time-dependent coherence was then estimated using a Morlet wavelet decomposition based on the method of Zhan et al (Zhan et al., 2006), with a resultant frequency resolution of 0.5 Hz and 95% confidence limits of 0.018.

#### Head motion predictability

Head motion predictability was quantified separately for linear head acceleration and angular head velocity according to a previously proposed procedure (MacNeilage & Glasauer, 2017). First, IMU signals were cut into individual strides synchronized to the left heel strike and further time-normalized by resampling each stride to a total of 300 samples. Head motion data from the first 125 strides (*N* = 125) was used for further analysis and averaged to reconstruct the mean head motion trajectory across the stride cycle, i.e., the stride-cycle attractor. Subsequently, the total variance *SS*_*tot*_ and residual variance *SS*_*res*_ of head motion were calculated:

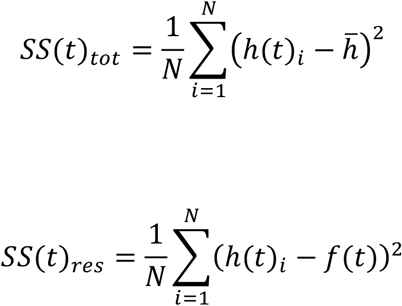

where *h*(*t*)_*i*_ is the head motion during the *i*th stride at the normalized stride time *t*, *h̃* is the average head motion over all stride cycle phases and strides, and *f*(*t*) denotes the stride cycle attractor. Correspondingly, *SS*_*tot*_ quantifies the signal deviation from the overall mean signal whereas *SS*_*res*_ gives the signal deviation from the stride cycle attractor.

Using these metrics, the proportion of head motion variance that can be explained by the stride cycle attractor, i.e., explained variance (*V*_*exp*_ = 1 − *SS*_*res*_/*SS*_*tot*_), and the proportion of residual head motion variance *V*_*res*_ = *SS*_*res*_/*SS*_*tot*_ can be derived. Low values of *V*_*res*_ indicate a high head motion predictability. Hence, knowing the exact stride cycle phase, feed-forward signals of the locomotor command can provide reliable information about the most likely ongoing movement. However, as *V*_*res*_ increases, head motion prediction based on stride cycle phase information becomes less accurate and additional sensory cues are required for head motion estimation. These considerations can be expressed in the form of a statistically optimal model, i.e., the maximum likelihood estimation model for cue integration (Ernst & Banks, 2002). Accordingly, head motion *Ĥ* can be estimated by a weighted linear combination of vestibular (sensory, *S*) and efference copy (motor, *M*) cues with weights *W*_*sens*_ and *W*_*mot*_ corresponding to the relative reliability of these cues:

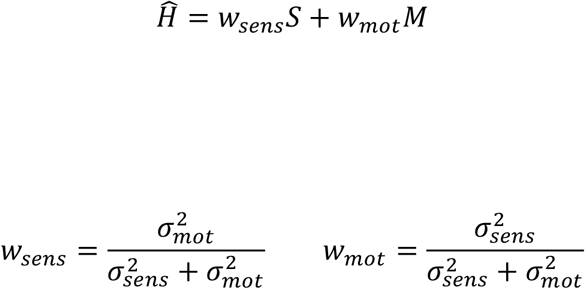

The above weights can now be estimated using the head motion data based on the following two assumptions: (1) According to Weber’s law, sensory noise is assumed to be signal-dependent, i.e., its variance should be proportional to the squared signal (Fechner, 1860). As the average signal is approximately zero for oscillatory locomotor movements, sensory noise can be estimated by 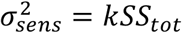 with the Weber’s fraction *k*. (2) If the intended head motion during each stride equals the stride cycle attractor, motor noise can be estimated as 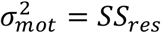 Based on these assumptions sensory weight can by expressed as directly proportional to *V*_*res*_:^1^

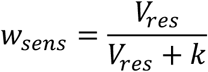

### 2.3 Statistical analysis

Data are reported as mean ± SD. The effects of correlation analysis parameters, head predictability estimates, and gait parameters were analyzed using a repeated-measures analysis of variance (rmANOVA) and Bonferroni post hoc analysis with condition (standing, slow and medium walking) as factor. Results were considered significant if *p* < 0.05. Statistical analysis was performed using SPSS (Version 25.0; IBM Corp., Armonk, NY).

## 3. Results

All participants exhibited significant correlations between SVS and COP displacements in both the frequency and time domain. SVS-COP coherence within the 0–25 Hz bandwidth peaked at 4.8 ± 1.4 Hz for standing, 3.5 ± 1.3 Hz for slow and 4.9 ± 1.4 Hz for medium walking speed (F_2,18_ = 2.4; p = 0.119; Figure 1A). Peak coherence dropped from standing to slow waking and further decreased with faster walking (F_2,18_ = 32.7; p < 0.001; Figure 1B). Cross-correlation analysis revealed a short latency component of SVS-induced COP responses at 85 ± 11 ms for standing and slightly later responses for slow (118 ± 22 ms) and medium (118 ± 15 ms) walking speeds (F_2,18_ = 15.5; p < 0.001). A medium latency response of opposite polarity occurred at 203 ± 45 ms for standing and slightly later for slow (274 ± 30 ms) and medium (284 ± 32 ms) walking speeds (F_2,18_ = 22.7; p < 0.001; Figure 1C). Phase lags at frequency bandwidth with significant coherence estimates corresponded to the medium latency response with 204 ± 44 ms for standing, and 270 ± 31 ms for slow, and 269 ± 35 ms for medium walking speed (F_2,18_ = 17.0; p < 0.001).

**Figure 1:**
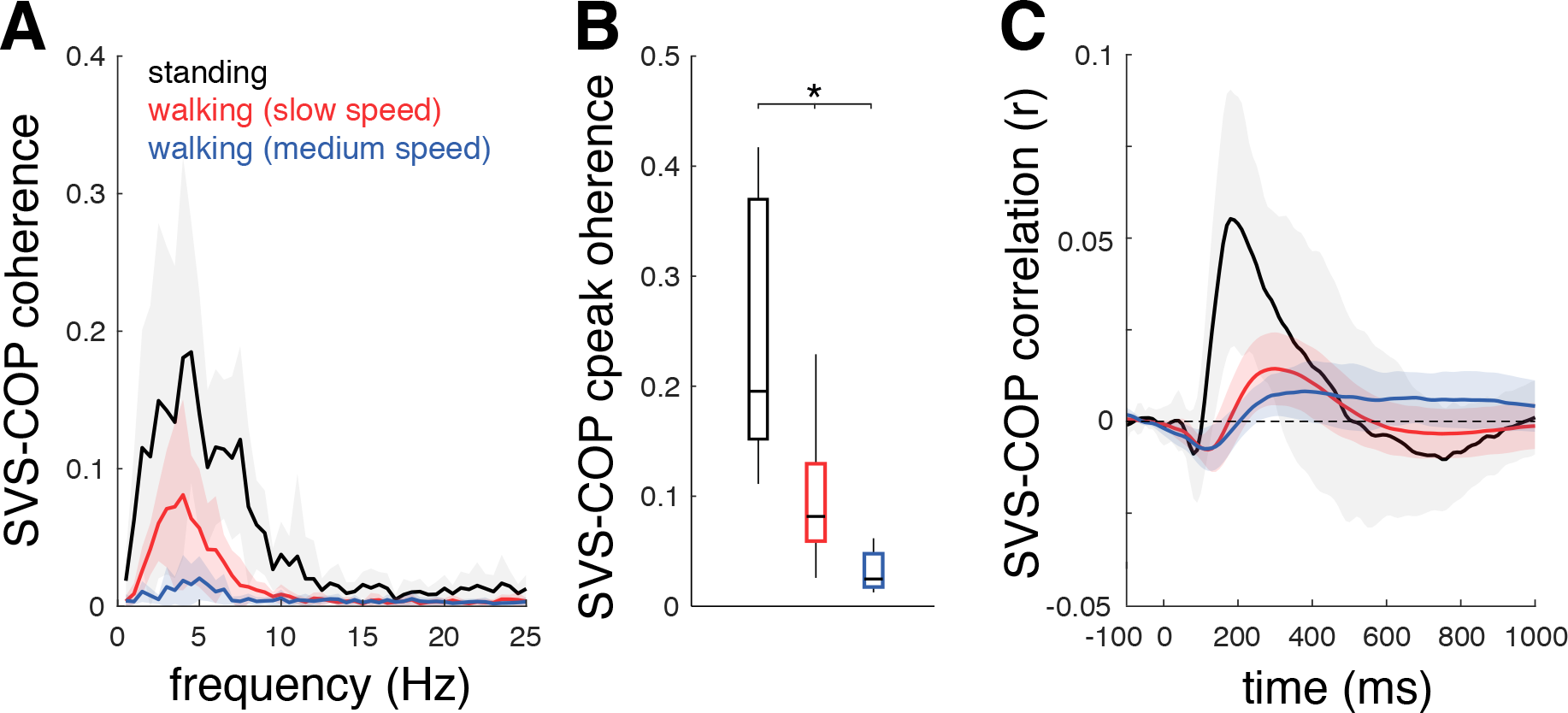
Correlation analysis in frequency and time domain for coupling between SVS and COP displacements during different activities. Coherence functions, (**B**) peak coherence values, and (**C**) corresponding cross-correlations between SVS and COP displacements. SVS-COP coherence drops from standing to slow walking and is further reduced at faster walking speed. SVS-induced COP displacements exhibit a short latency response around 80-120 ms and a medium latency response of opposite polarity at around 200-290 ms. * indicates a significant difference. *SVS: stochastic vestibular stimulation; COP: center-of-pressure*

Similar to global coherence estimates, time-frequency analysis of SVS-COP coupling across the gait cycle revealed a drop of peak coherence from slow to medium walking speed (F_1,9_ = 15.5; p < 0.001, Figure 2A-C). Analysis of head motion predictability revealed a corresponding decrease in mean head motion *V*_*res*_ (i.e., an increase in head motion predictability) from slow to medium walking speed for both linear head acceleration and angular head velocity (F_1,18_ = 14.0; p = 0.001, Figure 2A-C). Furthermore, head motion predictability was generally higher for linear head acceleration compared to angular head velocity (F_1,18_ = 44.7; p < 0.001, Figure 2C). SVS-COP coupling across the gait cycle exhibited phasic modulations with two distinct peaks occurring at 25.0 ± 2.4% and 74.3 ± 2.7% of the gait cycle during slow walking and at 25.4 ± 1.4% and 76.6 ± 1.4% of the gait cycle during walking at medium speed (Figure 2A,B). In accordance, the estimated head motion predictability was similarly modulated throughout the gait cycle. Periods of maximum *V*_*res*_ (i.e., least predictability) of angular head velocity corresponded to peaks of SVS-COP coherence (at 21.6 ± 2.8% and 73.5 ± 3.1% of the gait cycle during slow walking and 21.3 ± 1.3% and 73.4 ± 2.6% of the gait cycle during medium walking). In contrast, peaks of linear head acceleration *V*_*res*_ occurred at considerably earlier instances of the gait cycle (5.9 ± 2.8% and 56.4 ± 1.0% of the gait cycle during slow walking and 6.7 ± 2.9% and 56.9 ± 1.4% of the gait cycle during medium walking).

**Figure 2:**
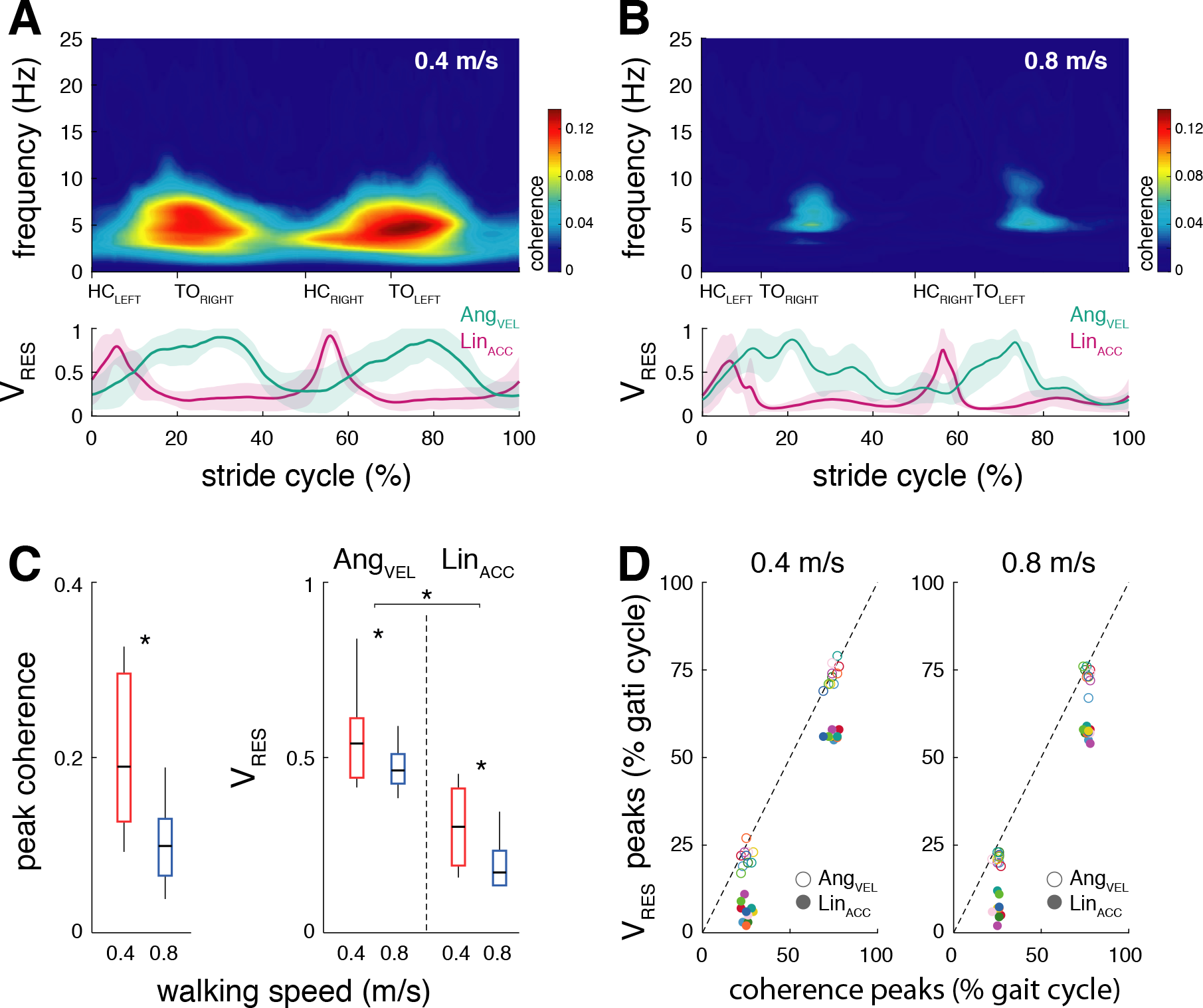
Time-frequency analysis of coupling between SVS and COP displacements and corresponding estimates of head motion predictability. (**A,B**) Average time-dependent coherence between SVS and COP at slow and medium walking speed (upper panels) and corresponding average head motion *V*_*res*_ curves (lower panels) in dependence on the gait cycle phase. (**C**) Peak coherence and corresponding average *V*_*res*_ for angular head velocity and linear head acceleration. (**D**) Temporal correspondence between phase-dependent peaks in SVS-COP coherence and peaks in head motion *V*_*res*_ at slow and medium walking speed. Both SVS-COP coupling as well as head motion predictability decrease with faster locomotion and are phase-dependently modulated across the gait cycle. SVS-COP coupling exhibits two peaks across the stride cycle that correspond well to periods of highest *V*_*res*_ (i.e., least predictability) of angular head velocity. * indicates a significant difference. *SVS: stochastic vestibular stimulation; COP: center-of-pressure; VRES: residual variance; AngVEL: angular head velocity; LinACC: linear head acceleration; HC: heel contact; TO: toe off*

Continuous SVS did not affect the average spatiotemporal walking pattern but resulted in a considerable increase of stride-to-stride variability (i.e., increased CV) of all analyzed gait parameters (Figure 3B). This effect was diminished during medium compared to slow walking for the CV of stride time (F_1,9_ = 5.9; p = 0.038), single support percentage (F_1,9_ = 7.7; p = 0.022), and double support percentage (F_1,9_ = 5.9; p = 0.038).

**Figure 3:**
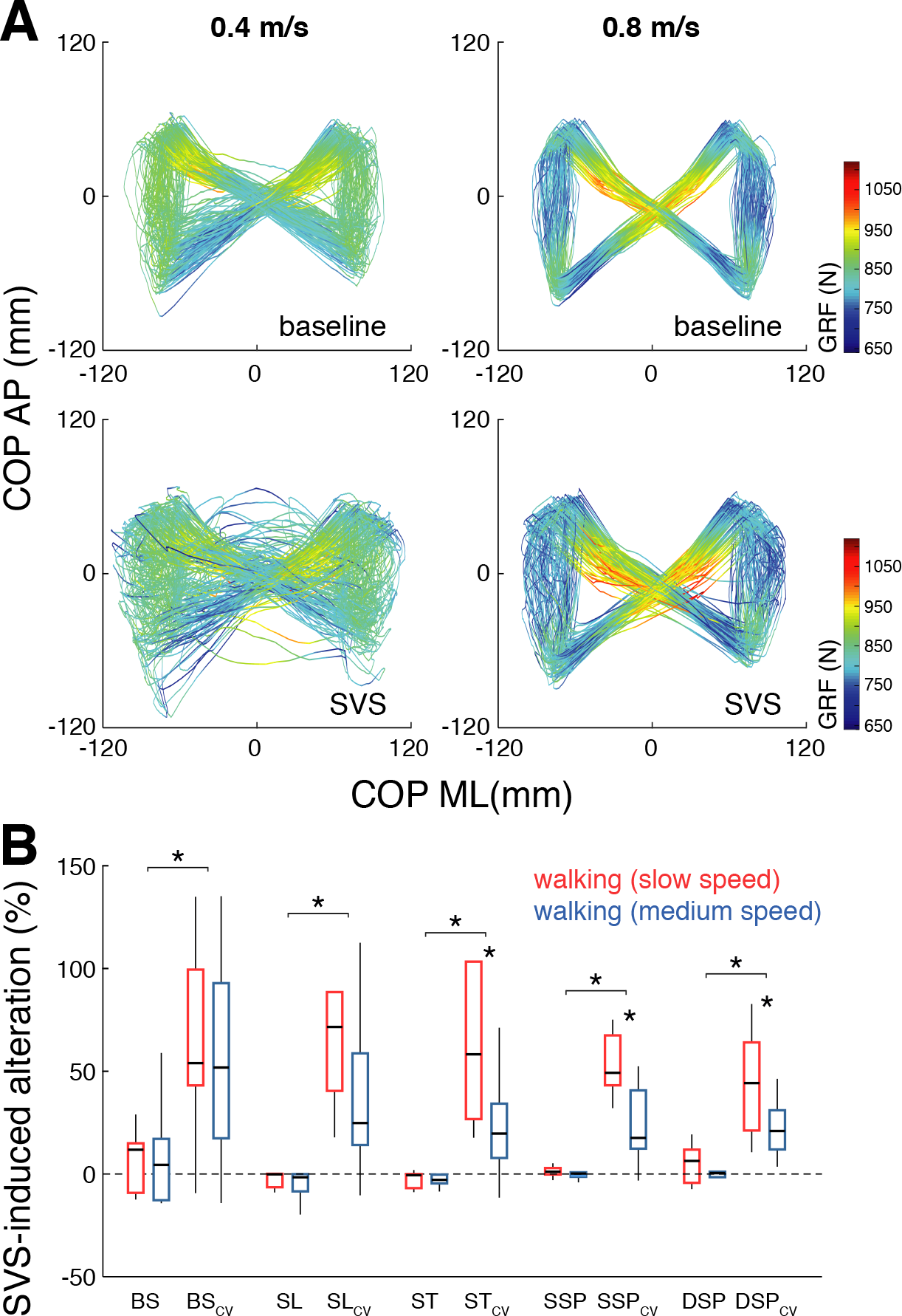
Effects of SVS and walking speed on spatiotemporal gait parameters. (**A**) Representative examples of COP trajectories during slow (left) and medium (right) walking speed for trials without stimulation (upper panel) and with continuous SVS (lower panel). (**B**) Percentage differences in gait parameters for walking with continuous SVS compared to baseline walking at the two locomotor speeds. SVS does not affect the mean gait pattern but induces increased stride-to-stride fluctuations (i.e., increased CV values) in all gait parameters. This effect diminishes with faster locomotion. * indicates a significant difference. *SVS: stochastic vestibular stimulation; COP: center-of-pressure; BS: base width; SL: stride length; ST: stride time; SSP: single support percentage; DSP: double support percentage; CV: coefficient of variation*

## 4. Discussion

Here we observed that activity-dependent modulations of vestibular influence on balance control closely match differences in head motion predictability. This finding supports a previously proposed model (MacNeilage & Glasauer, 2017), based on the idea that during stereotyped locomotion, efference copies of locomotor commands may be used in conjunction with sensory, especially vestibular, cues in order to estimate resultant head movements and trigger adequate balance adjustments. The extent to which balance regulation during locomotion relies on concurrent vestibular vs. motor feed-forward signals should further depend on the reliability of these estimates, such that higher weighting is given to the less noisy estimate (Ernst & Banks, 2002). Accordingly, we found that activities linked to less stereotyped head movements (i.e., standing or slow walking) were more sensitive to externally triggered vestibular cues than activities with highly predictable head motion patterns (i.e. faster walking). Furthermore, we found that during walking, sensitivity to SVS was highest at the times of lowest head movement predictability. Thus, the present results provide a reasonable explanation for the dynamic weighting of vestibular influences across and within different activities and further emphasize the possibility of an intrinsic feed-forward regulation of balance during human locomotion based on locomotor efference copies. In the following, we will discuss these findings with respect to their functional implications and possible physiological correlates.

The influence of externally triggered vestibular cues on body sway (i.e., SVS-COP coherence) was attenuated during walking compared to standing (Figure 1). This agrees with the recently reported decrease of vestibular influence on body balance after gait initiation and the corresponding increase after gait termination (Tisserand et al., 2018). Such general down-weighting of vestibular influence during locomotion is consistent with predictions of the model employed here. During locomotion, the presence of an efference copy of locomotor commands imposes an upper limit for the weighting of sensory influences, i.e., *W*_*sens*_ < 1/(1 + *k*) < 1, which depends on the Weber’s fraction *k*, the proportionality constant for signal dependent noise (MacNeilage & Glasauer, 2017). Thus, in contrast to standing, balance regulation during locomotion will always be partially governed by a locomotor efference copy, i.e., *W*_*mot*_ > 0. Previous literature indicates that the attenuation of balance-related vestibular reflex gains during locomotion is a more general phenomenon that also concerns vestibulo-ocular reflex pathways (Dietrich & Wuehr, 2019). Accordingly, it was shown in patients with a unilateral vestibular failure that spontaneous nystagmus resulting from a vestibular tone imbalance is considerably dampened during ambulation (Jahn et al., 2002). A complete suppression of the horizontal vestibulo-ocular reflex has been demonstrated in tadpole swimming, i.e., a locomotor activity where the spatiotemporal coupling between rhythmic propulsive locomotor movements and resultant head displacements is high (Chagnaud et al., 2012; Lambert et al., 2012). Similar effects were also observed in other non-vestibular sensory modalities. For instance, proprioceptive stretch reflexes that govern postural control during standing are known to be selectively suppressed during locomotion (Dietz et al., 1985).

During locomotion, vestibular feedback is thought to be essential for maintenance of dynamic stability by fine-tuning the timing and magnitude of foot placement (Bent et al., 2004; Blouin et al., 2011; Wuehr et al., 2016). In line with this, significant SVS-COP coupling during locomotion led to an increased spatiotemporal variability of stride-to-stride walking movements despite of the otherwise unaffected average gait parameters (Figure 3). Both SVS-COP coupling and increased stride-to-stride variability decreased from slow to medium walking speed (Figure 1-3). This observation is in line with previous studies reporting that the destabilizing impact of a vestibular loss or external vestibular perturbation is considerably attenuated during fast compared to slow locomotion (Brandt et al., 1999; Dakin et al., 2013; Jahn et al., 2000; Schniepp et al., 2012). Moreover, in agreement with previous reports (Bent et al., 2004; Blouin et al., 2011; Dakin et al., 2013), we found that SVS-COP coupling was phase-dependently modulated during locomotion, exhibiting two consistent peaks across the gait cycle with equal timing for both examined walking speeds. Both speed- and phase-dependent changes in SVS-COP coupling closely matched concomitant changes in head motion predictability (Figure 2). Accordingly, *V*_*res*_ of linear acceleration and angular velocity of head motion decreased with faster locomotion (i.e., increased predictability) and consistently exhibited two local maxima across the gait cycle (i.e., least predictability). Furthermore, phasic modulation of SVS-COP coupling across the locomotor cycle temporally matched modulations of *V*_*res*_ of angular head velocity rather than of linear head acceleration. This suggests that the observed SVS-induced COP displacements primarily reflect responses to activation of semicircular canal afferents conveyed through vestibulospinal tracts. In line with this, semicircular canal afferent stimulation was previously shown to trigger medium-latency body sway responses at the frequency bandwidth and phase lags observed in the present study, whereas short-latency responses triggered by otolith afferents via reticulospinal tracts occur at a higher bandwidth (> 10 Hz) with shorter phase lags (Cathers et al., 2005; Dakin et al., 2007).

Previous reports hypothesized that the activity-dependent modulation of vestibular feedback during locomotion is reflected by concurrent changes in muscle activation or foot placement patterns to stabilize posture (Bent et al., 2004; Blouin et al., 2011; Dakin et al., 2013). Others have proposed that vestibular down-regulation during faster locomotion simply occurs due to a larger degree of automated behavior that tends to rely less on sensory feedback (Brandt et al., 1999). The present findings alternatively suggest that rather than automation, changes in head motion predictability define the activity-depended modulation of vestibular control of balance during locomotion. Accordingly, the ratio of sensory vs. motor noise generally becomes greater with increasing locomotion speed, which should lead to a down-weighting of vestibular feedback in favor of a direct feed-forward regulation of balance based on efference copies from the locomotor commands. Moreover, the phase-dependent modulation of vestibular influences would similarly reflect changes in the proportion of sensory vs. motor noise across the gait cycle. An analogous re-weighting of sensory vs. motor cues based on the relative precision of these signals could further explain the previously described speed- and phase-dependent modulation of other non-vestibular feedback cues (i.e., visual and proprioception) occurring during human locomotion (Dietz, 2002; Jahn et al., 2001; Logan et al., 2014; Wuehr et al., 2013; Wuehr et al., 2014).

The relationship between the activity-dependent modulation of vestibular influences and changes in head motion predictability suggests that during human locomotion an intrinsic feed-forward mechanism based on locomotor efference copies plays a part in balance regulation, which was previously thought to be purely controlled by sensorimotor reflexes. Traditionally, motor efference copies are primarily considered to serve as predictors of sensory consequences arising from one’s own actions, thereby enabling the brain to distinguish self-generated sensory signals (reafference) from sensory inputs caused by unpredictable external influences (exafference) (Cullen, 2004; Sperry, 1950; von Holst & Mittelstaedt, 1950). Recent research, however, has expanded this view, suggesting that internal motor predictions are also involved in coordinating action of different motor systems that are otherwise functionally and anatomically unrelated (Straka et al., 2018). One well described example of such an efference copy-mediated motor-to-motor coupling is the interaction between the mammalian locomotor and respiratory motor system, which is coordinated by intrinsic efference copies derived from CPG activity in the lumbar spinal cord (Onimaru & Homma, 2003). More recently, CPG-derived locomotor efference copies were shown to directly mediate compensatory eye movements for gaze stabilization during aquatic locomotion in *Xenopus laevis* tadpoles (Lambert et al., 2012) and adult frogs (von Uckermann et al., 2013) – a task that is usually thought to be mediated by the vestibulo-ocular reflex. Moreover, this direct coupling between spinal and ocular motor signals was shown to be accompanied by a selective suppression of vestibular inputs to extraocular motoneurons. Whether such selective gating of vestibular feedback occurs at the level of the brainstem extraocular or vestibular nuclei or other brain regions such as the cerebellum yet remains unknown. In favor of a cerebellar origin, it was previously shown that the phasic modulation of vestibulospinal neuron activity in the lateral vestibular nucleus observed during locomotion in cats depends on the presence of an intact cerebellum and is disrupted by its removal (Orlovsky, 1972; Udo et al., 1982). Given its prominent role in adaptive plasticity of vestibular reflexes (Angelaki & Cullen, 2008; Dietrich & Straka, 2016; Gittis & du Lac, 2006), the cerebellum might thus serve as a convergence site for the weighting and integration of self-motion derived vestibular cues and intrinsic locomotor efference copies.

## Acknowledgements

This work was supported by the German Federal Ministry of Education and Research (01EO1401) and the German Research Foundation (MA 6233/1-1).

## Footnotes

1 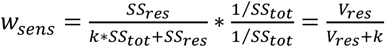

